# An integrated ‘omics approach highlights the role of epigenetic events to explain and predict response to neoadjuvant chemotherapy and bevacizumab

**DOI:** 10.1101/2022.07.06.498803

**Authors:** Thomas Fleischer, Mads Haugland Haugen, Jørgen Ankill, Laxmi Silwal-Pandit, Anne-Lise Børresen-Dale, Ingrid Hedenfalk, Thomas Hatschek, Jörg Tost, Olav Engebraaten, Vessela N. Kristensen

## Abstract

Here we present an integrated ‘omics approach for DNA methylation profiling using copy number alteration, gene expression and proteomic data to predict response to therapy and to pinpoint response-related epigenetic events. Fresh frozen tumor biopsies taken before, during and after treatment from patients receiving neoadjuvant chemotherapy with or without the anti-angiogenic drug bevacizumab were subjected to molecular profiling. Our previous studies have shown that administration of bevacizumab in addition to chemotherapy (combination treatment) may confer improved response for patients; here we report that DNA methylation at enhancer CpGs related to cell cycle regulation can predict response to chemotherapy and bevacizumab for ER positive patients with high fidelity (AUC=0.874), and we validate this observation in an independent patient cohort with similar treatment regimen (AUC=0.762). When combining the DNA methylation score with a previously reported proteomic score (ViRP), the prediction accuracy further improved in the validation cohort (AUC=0.784). We also show that tumors receiving the combination treatment underwent more extensive epigenetic alterations than tumors receiving only chemotherapy. Finally, we performed an integrative emQTL analysis on alterations in DNA methylation and gene expression levels, showing that the epigenetic alterations that occur during treatment are different between responders and non-responders and that these differences may be explained by the proliferation-EMT axis through the activity of the transcription factor GRHL2. Taken together, these results illustrate the clinical benefit of the addition of bevacizumab to chemotherapy if administered to the correct patients.

## Introduction

Neoadjuvant treatment is commonly used for patients with large breast tumors to avoid delays in important systemic therapy, to evaluate the efficacy of such therapy and facilitate the surgical resection of breast tumors. With the rationale of denying tumors access to nutrients and oxygen through vascular endothelial growth, anti-angiogenic therapy has been applied to decrease tumor size in neoadjuvant setting [1-3]. Bevacizumab is a monoclonal antibody that binds to vascular endothelial growth factor (VEGF) and blocks the binding to its receptors thus inhibiting endothelial growth [4]. Although the drug has shown efficacy in some trials [5-10], it did not increase overall survival. We have previously shown that Bevacizumab can improve response to neoadjuvant chemotherapy in patients with ER positive tumors [11]. A better understanding of the biological mechanisms leading to response or resistance to this treatment as well as biomarkers for prediction of response and benefit of adding bevacizumab to chemotherapy are needed. Recently, a signature consisting of the expression of nine proteins was shown to predict response to the combination of chemotherapy and bevacizumab in the NeoAva trial with an area under the ROC curve (AUC) of 0.92 for ER positive and 0.85 for all patients [12]. A methylation signature, predictive of response to chemotherapy and bevacizumab has been reported with an AUC of 0.91 in a study with patients with triple negative breast tumors [13], and a hypoxia gene score has been associated with response to chemotherapy and bevacizumab in patients with HER2 negative tumors (OR=2.4) [14]. Together, these reports highlight the potential for use of molecular signatures for prediction of response.

We have previously shown that DNA methylation of cell cycle genes changes during treatment with chemotherapy (Doxorubicin or Fluorouracil (5FU) and mitomycin C), and that these changes differ between responders and non-responders [15]. To study the biological significance of epigenetic changes, we integrated DNA methylation and mRNA expression using an expression-methylation quantitative trait loci (emQTL) analysis. The emQTL approach enabled us to identify variation in DNA methylation with concomitant variation in gene expression in breast tumors. This led to the discovery of a mechanism to maintain continuously active estrogen receptor signaling in ER positive breast tumors through loss of DNA methylation at enhancers carrying binding sites for the transcription factors (TFs) ER, FOXA1 and GATA3 [16], and loss of methylation at enhancers carrying binding sites of TFs related to proliferation in ER negative tumors [17]. Treatment-induced epigenetic changes at enhancers through DNA methylation have to our knowledge not yet been investigated.

In the current study, we aimed at developing a predictive signature based on DNA methylation that could assist in selecting patients for the combination therapy of chemotherapy and bevacizumab. We also investigated alterations in DNA methylation that were both common to all patients as well as patient-specific and could explain varying response to treatment. Here, we report that DNA methylation at enhancer CpGs can predict response to the treatment with chemotherapy and bevacizumab for ER positive patients, and we present evidence that tumors receiving the combination treatment underwent more extensive overall epigenetic alterations during treatment than tumors receiving only chemotherapy. Also, the epigenetic alterations occurring during treatment differ between responders and non-responders and these differences may be explained by the proliferation-EMT axis through the activity of the transcription factor GRHL2.

## Materials and methods

### Study design and patient material

The NeoAva clinical trial has been reported previously [11]. In brief, patients were recruited at two hospitals in Norway between 2008 and 2012, and written informed consent was obtained from all patients prior to inclusion. The study was approved by the Institutional Protocol Review Board, the regional ethics committee, the Norwegian Medicines Agency, and carried out in accordance with the Declaration of Helsinki, International Conference on Harmony/Good Clinical practice. The study is registered in the ClinicalTrials database with the identifier NCT00773695. Patients with HER2-negative mammary carcinomas with size >2.5 cm previously untreated for breast cancer and no sign of metastatic disease were eligible. The patients were randomized 1:1 to receive bevacizumab and chemotherapy (combination treatment arm) or chemotherapy alone (chemotherapy arm).

The chemotherapy regimen consisted of four cycles of FEC100 (5-fluorouracil 600 mg/m2, epirubicin 100 mg/m2, and cyclophosphamide 600 mg/m2) every 3 weeks, followed by docetaxel 100 mg/m2 every 3 weeks or 12 weekly infusions of paclitaxel 80 mg/m2. Bevacizumab was administered intravenously at a dose of 15 mg/kg every third week or 10 mg/kg every other week in patients receiving docetaxel or paclitaxel, respectively. Samples were sequentially collected before treatment (core needle biopsies; termed “week 0” samples), during treatment (core needle biopsies 12 weeks into treatment; termed “week 12” samples), and after treatment (at surgery; termed “week 25” samples). Response to treatment was measured as either pathologic complete response (pCR; complete eradication of all invasive cancer cells in both breast and axillary lymph nodes), residual cancer burden (RCB; [18]), and as fraction of tumor volume left after treatment.

### DNA extraction and DNA methylation profiling

Fresh frozen tumor biopsies were dissected into small pieces, mixed, and divided into amounts suitable for DNA, RNA, and protein extraction. DNA was isolated using the QIAcube and AllPrep DNA/RNA Mini Kit 350 or 600 for biopsies from the first two or the last time point, respectively (Qiagen) according to the company’s standard protocol. The DNA methylation status of more than 450,000 CpG sites was interrogated using the Infinium HumanMethylation450 microarray (Illumina). Preprocessing and normalization involved steps of probe filtering, color bias correction, background subtraction and subset quantile normalization as previously described [19]. Raw and normalized data is available from the Gene Expression Omnibus (GEO) with accession number GSE207460.

### mRNA and copy number profiling

Gene expression profiling has been previously reported [11]. Briefly, the analysis was performed using one color SurePrint G3 Human GE 8×60k Microarrays (Agilent Technologies), and the data is available in the ArrayExpress database (http://www.ebi.ac.uk/arrayexpress) under accession number E-MTAB-4439. The PAM50 subtyping algorithm [20] was used to determine the molecular subtype of each sample.

Copy number alteration (CNA) profiles has previously been reported [21]. Briefly, tumor DNA was analyzed for CNAs using Genome-Wide Human SNP array 6.0 (Affymetrix). The copy number profiles were segmented with the allele-specific piecewise constant fitting (ASPCF) algorithm [22], and subsequently, allele-specific copy number analysis of tumors (ASCAT; [23]) was used to estimate tumor cell fraction, tumor ploidy, and copy number. The data is deposited at the European Genome-phenome Archive under accession number EGAS00001003287.

### Validation data

Validation for the predictive signature for response to chemotherapy was performed in the Promix trial [24], a single arm phase II neoadjuvant trial where patients with HER2 negative tumors received chemotherapy in combination with bevacizumab. Gene expression data is available through GEO with accession number GSE87455. DNA methylation data was not available for this patient cohort.

Validation for the concomitant alterations in DNA methylation and gene expression was performed in the Doxo/Fumi data set [15, 25, 26]. In this study, DNA methylation was measured for paired tumor samples from locally advanced breast cancer patients treated with doxorubicin and Fluorouracil (5FU) and mitomycin C.

### Statistical analyses

All statistical and bioinformatical analyses were carried out in the R software version 4.1.2 [27] unless otherwise specified.

### Prediction of response to combination treatment

Prediction of patient achieving pCR from DNA methylation data was performed using the machine learning method LASSO [28]. This method is susceptible to overfitting if the number of features is too large and if there are features with no biological importance. We thus performed a preselection of CpGs, restricting the data to include only CpGs indicated to be involved in epigenetic regulation of processes considered important in primary breast tumors based on our recently reported improved emQTL analysis [17]. We identified 44,263 CpGs involved in four mechanisms related to breast tumor biology. A CpG was selected when either involved in 1) estrogen-regulated proliferation in tumor cells, 2) non-estrogen regulated proliferation in tumor cells, 3) varying immune infiltration, or 4) varying fibroblast infiltration. These 44,263 CpGs were used in the LASSO analysis.

We constructed Lasso models using the R package *glmnet* [29] in an internal validation strategy (leave-one-out; LOO) where models are trained on N-1 samples (where N is the total number of samples in the analysis) and applied to the one remaining sample. This is run N times to construct N models and assign all samples a probability of achieving pCR. To avoid selection bias we also performed nested cross-validation (CV; function *cv.glmnet*), where another 5-fold CV was applied within each sub-training set to determine the optimal model parameters, e.g., the weight and beta values of the Lasso model. The other model parameters were set to their default values. Receiver operating characteristic (ROC) curves were generated using the *pROC* R package [30].

To translate a DNA methylation signature to a gene expression signature, we selected the genes whose expression had the highest absolute correlation to the CpGs in the signature. The beta coefficients of the linear model to be applied on gene expression data were determined by multiplying the beta coefficients for each CpG of the linear model of DNA methylation with the correlation coefficients (rho values) between DNA methylation and gene expression.

### Hybrid protein-methylation prediction score

To compare and improve the performance of the DNA methylation-based and protein-based prediction of response to the combination therapy, we generated a hybrid score integrating the previously published nine-protein ViRP score [12] and the discovered DNA methylation score. In the validation data set (Promix [24]), both signatures were translated to mRNA signatures, the scores were standardized (mean=0; sd=1), and the average value was calculated for each patient.

### Identification of treatment induced changes in DNA methylation

To assess treatment-induced changes in DNA methylation, we performed paired t-tests (R function *t.test*) between week 0 samples and week 12, as well as between week 12 samples and week 25 samples. CpGs were included in the analysis if the inter-quartile range (IQR) was larger than 0.1. Alterations were considered significantly different if the Benjamini-Hochberg corrected p-value was smaller than 0.05 and the absolute change in DNA methylation was larger than 0.1.

### Identification of concomitant changes in DNA methylation and gene expression (delta emQTL)

To identify epigenetic alterations with concomitant changes in genes expression, potentially highlighting transcriptional programs under epigenetic control that are altered during treatment, we performed an emQTL analysis [16, 17] using delta methylation and delta expression levels (changes between week 0 and week 12). Correlation analyses were performed between all possible CpG-gene pairs, and correlations were considered significant if the Bonferroni-corrected p-value was less than 0.05. Bonferroni correction of p-values in this analysis was chosen to conform to previous reports [16, 17].

For all CpGs and genes with at least one significant correlation, spectral co-clustering (biclustering) was performed on the inverse correlation coefficients using the *SpectralCoclustering* algorithm (random state=0) contained within the scikit-learn library [31] in Python version 3.7.9.

### Gene set enrichment analysis and hierarchical clustering

Gene sets used for gene set enrichment analysis were downloaded from the Molecular Signatures Database v7.4 (MSigDB) [32]. Enrichment was determined by hypergeometric testing (R function *phyper*) using the Hallmark (H) and gene ontology (GO; C5) gene set collections. P-values were corrected for multiple testing using Benjamini-Hochberg procedure (R function *p.adjust*).

Hierarchical clustering of the DNA methylation and gene expression levels was performed using the R package *pheatmap* [33] using Euclidean distance and ward.D2 linkage.

### Genomic segmentation and annotation

ChromHMM segmentation data from cell lines representing different breast cancer subtypes were obtained from Xi et al. [34], and included MCF7 and ZR751 (Luminal A), UACC812 and MB361 (Luminal B), HCC1954 and AU565 (HER2+), HCC1937 and MB469 (Basal-like). In this work, ChIP-seq peaks from key histone modifications including H3K4me3, H3K4me1, H3K27me3, H3K9me3 and H3K36me3 were used to predict chromatin states across the genome of the cell lines. The genomes were annotated into thirteen distinct chromatin states including: active promoter (PrAct), active promoter flanking (PrFlk), active transcription (TxAct), active transcription flanking (TxFlk), active intergenic enhancer (EhAct), active genic enhancer (EhGen), bivalent promoter (PrBiv), bivalent enhancer (EhBiv), repressive polycomb domain (RepPC), weak repressive domain (WkRep), repeat/ ZNF genes (RpZNF), heterochromatin (Htchr) and quiescent state / low signals (QsLow) [34]. Subtype specific ChromHMM annotations were made by collapsing the ChromHMM annotations from cell lines of similar subtype, omitting genomic regions with discordant annotation within a subtype. Enrichment of CpGs in a ChromHMM defined functional region was measured as the ratio between the frequency of delta emQTL CpGs found in a specific segment type over the frequency of CpGs from the Illumina HumanMethylation450 array found within the same segment type. P-values were obtained by hypergeometric testing with the Illumina 450k array probes as background (n=485,512). P-values were corrected for multiple testing using Benjamini-Hochberg procedure.

### TF enrichment analysis in UniBind defined TF binding sites

Enrichment of CpGs in TF binding sites was assessed using data obtained from the UniBind database [35]. Maps of direct TF-DNA interactions were downloaded from the UniBind website (https://unibind2018.uio.no). The genomic positions of all CpGs from the Illumina 450k array lifted over from hg19 to hg38 using the LiftOver webtool from UCSC genome browser (https://genome.ucsc.edu) and extended with 150 bp upstream and downstream. Since each TF can have multiple TF binding sites derived from different ChIP-seq experiments, we merged the TF binding sites for all ChIP-seq experiments for each TF. Enrichment of CpGs in TF binding sites was computed using hypergeometric testing (R function *phyper*) with IlluminaMethylation450 Bead Chip CpGs as background. False discovery rate was estimated by Benjamini-Hochberg correction.

### Chromatin interaction mapping

Chromatin Interaction Analysis by Paired-End Tag sequencing (ChIA-PET) is a method to identify promoter-enhancer loops on a genome-wide scale [36]. ChIA-PET data defining long-range chromatin interactions in the ER+ MCF7 breast cancer cell line was obtained from ENCODE (Accession number ENCR000CAA; [36]). Only *cis* loops were included in the analysis. An emQTL was considered to be in a ChIA-PET Pol2 loop if the CpG and transcription start site of its associated gene were found within the genomic intervals of two opposite ends of the same loop. Enrichment of *in cis* (i.e. on the same chromosome) emQTLs in ChIA-PET loops was determined by hypergeometric tests (R function phyper) using all possible *in cis* CpG-gene pairs as background.

### Generation of tumor purity corrected DNA methylation profiles

To determine tumor cell-specific alterations in DNA methylation, we developed a method to estimate the tumor cell-specific methylation level in biopsies with varying tumor purity. To achieve this, we utilized ASCAT-estimated tumor purity [23] generated from CNA measurements in the same tumors [21]. For each CpG, a linear regression was constructed between the ASCAT-estimated tumor purity and measured DNA methylation. The methylation level for the regression line at tumor purity 100% was considered the mean tumor cell methylation, and each sample’s distance to the regression line (residual) added to the mean tumor cell methylation was made to represent the tumor methylation. Consequently, each CpG was adjusted independently, depending on its association to tumor purity. This approach allowed us to analyze two versions of the DNA methylation data: original DNA methylation and tumor purity-adjusted DNA methylation.

## Results

DNA methylation profiling was performed on fresh frozen core biopsies from breast tumors from patients receiving neoadjuvant chemotherapy with or without bevacizumab. DNA methylation profiling was performed on DNA from biopsies taken before treatment (week 0), after FEC treatment (weeks 12), and surgical specimens after taxane treatment (week 25).

### DNA methylation in pre-treatment primary tumors predicts response to combination therapy with chemotherapy and bevacizumab

To assess the potential of DNA methylation in pre-treatment tumor samples to predict treatment response, we performed a machine learning approach (LASSO) to identify the CpGs most predictive of pathologic complete response (pCR). 44,263 CpGs identified in our previous studies [17] to be involved in epigenetic de-regulation (and associated to gene expression) in one of four important functions in breast cancer were used in the LASSO analysis: 1) estrogen signaling, 2) non-estrogen regulated proliferation in tumor cells, 3) varying immune infiltration, or 4) varying fibroblast infiltration were further studied for their potential to predict treatment response.

To produce a predictive signature and avoid overfitting, we performed an internal validation approach (leave-one-out cross-validation) and executed the analysis on each treatment arm individually to avoid mixing the response to two different treatment regimens. Further, the analysis was performed on ER positive and ER negative samples separately, or independently of ER status. For patients in the combination arm with ER positive tumors and available DNA methylation profiles (n=53), the leave-one-out approach was able to identify patient who would achieve pCR using pre-treatment samples, giving an area under the ROC curve of 0.874 (**Figure 1A**). After selection of an optimal cutoff, we obtained a sensitivity of 82% and a specificity of 95%. To obtain a stable predictive signature, we combined the 53 models by averaging the linear regression beta coefficients for CpGs present in more than 50% of the models yielding an 11 CpG signature (**Table 1**). This stable model was applied to the DNA methylation data, perfectly predicting patients achieving pCR (area under the ROC curve of 1; **Supplementary Figure 1A**).

**Figure 1:**
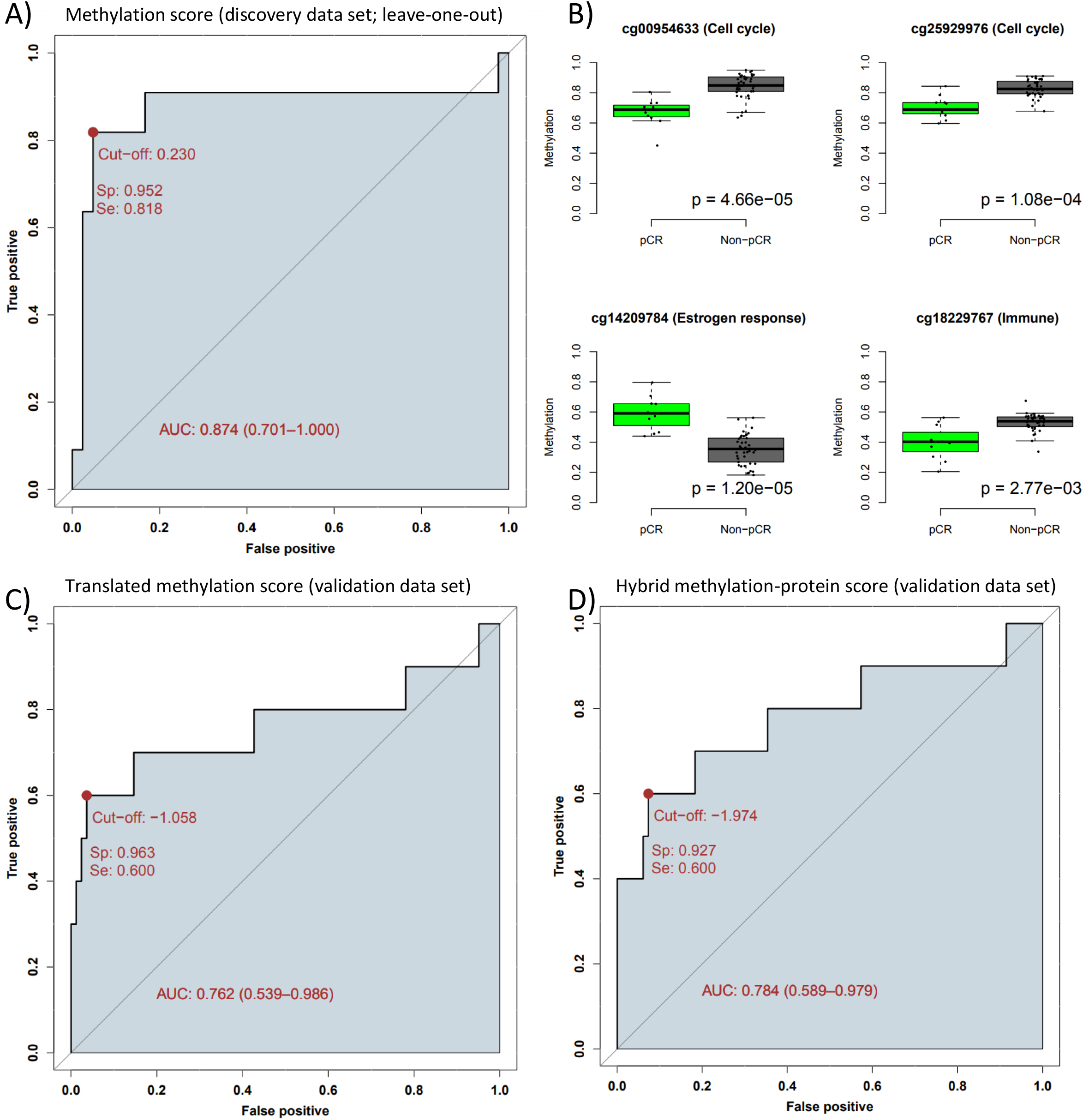
Identification of predictive signatures for response to neoadjuvant treatment with chemotherapy plus bevacizumab in ER positive breast cancer. A) ROC curve showing specificity vs sensitivity for the leave-one-out cross-validation probabilities for response. B) Boxplots of DNA methylation levels in pre-treated samples for example CpGs in the identified predictive signature. P-value is determined using t-test. C-D) ROC curves showing specificity vs sensitivity in the validation cohort C) the “translated” gene expression signature, and D) the hybrid protein-methylation score. AUC: area under the curve.

**Table 1:** Predictive signatures for response to neoadjuvant treatment with chemotherapy plus bevacizumab in ER positive breast cancer. Aggregate predictive signature for response to therapy. Beta methylation is the linear regression coefficient for DNA methylation, Gene is the gene with highest absolute correlation to the CpG, Correlation meth-expr is the Pearson correlation coefficient between DNA methylation and gene expression, Beta expression is the “translated” linear regression coefficient (i.e., Beta methylation multiplied by Correlation meth-expr), emQTL cluster is the previously identified biological function of the CpG.

| Probe | Beta methylation | Gene | Correlation meth-expr | Beta expression | emQTL cluster |
| --- | --- | --- | --- | --- | --- |
| cg00954633 | -0.42 | PALM | 0.64 | -0.27 | Cell cycle |
| cg25929976 | -0.20 | CTSC | -0.52 | 0.10 | Cell cycle |
| cg19901005 | -0.18 | SH2D2A | -0.58 | 0.10 | Cell cycle |
| cg12445422 | -0.16 | YBX1 | -0.67 | 0.10 | Cell cycle |
| cg16004056 | -0.09 | TBX19 | -0.68 | 0.06 | Cell cycle |
| cg12772314 | -0.07 | ATP6AP1L | 0.41 | -0.03 | Cell cycle |
| cg14209784 | 0.97 | TNFAIP3 | 0.64 | 0.62 | Estrogen response 1 |
| cg05870586 | 0.39 | GALNT10 | -0.56 | -0.22 | Estrogen response 1 |
| cg08287887 | 0.08 | JMJD8 | -0.60 | -0.05 | Estrogen response 1 |
| cg18229767 | -0.09 | TNFSF14 | -0.60 | 0.06 | Immune |
| cg09524658 | -0.08 | MLKL | -0.62 | 0.05 | Immune |

The majority of the CpGs in the predictive pre-treatment signature were associated with cell cycle regulation, three CpGs were associated with estrogen response, and two CpGs were associated with varying immune infiltration. The CpGs were significantly differently methylated between responders and non-responders (examples in **Figure 1B**), with CpGs involved in cell cycle regulation and immune infiltration being less methylated in responders, and CpGs involved in estrogen response being more methylated in responders.

The fact that the pre-selected CpGs were also strongly associated to expression levels of genes [17] enabled us to calculate a “translated” signature by multiplying the methylation beta coefficients with the correlation coefficients of the gene with highest absolute correlation, thus constructing a predictive signature that could be applied also on gene expression data (**Table 1**). When applying this signature to the gene expression data (**Supplementary Figure1B**) we obtained an area under the ROC curves of 0.870. We then performed validation of the predictive signature in an independent cohort: the Promix clinical trial [24]. Since no DNA methylation data was available for this cohort, we applied the predictive “translated” gene expression signature and observed an area under the ROC curve of 0.762 and with optimal cutoff selection we obtained a sensitivity of 60% and a specificity of 96% in the validation expression dataset (**Figure 1C**).

When performing the analysis on both ER positive and ER negative tumors, the identified signature from each iteration of the LOO approach was different, and no stable model could be determined; also, the prediction was poorer (data not shown). Results with translational value could not be generated from the chemotherapy arm (too few patients with pCR), or only ER negative patients in the combination arm (too few patients). Also, when applying the methodology directly on gene expression data, the prediction performance was inferior in both discovery and validation data.

### A hybrid protein and DNA methylation score improves prediction of treatment response from pre-treatment biopsies

To assess the link between the identified DNA methylation signature and the published ViRP protein signature [12] we calculated the correlation between the scores. The scores were highly correlated (Pearson correlation coefficient of 0.71; p-value << 0.05; **Supplementary Figure 1C**). Further, to assess the complementarity of the two scores, we generated a hybrid score by averaging the scores from the two signatures. When applying the hybrid score on the validation set (Promix), we obtained an area under the ROC curve of 0.784; selecting the optimal cutoff-value we obtained a sensitivity of 60% and a specificity of 93% (**Figure 1D**).

### DNA methylation profiles after treatment with chemotherapy +/- Bevacizumab

To identify alterations in DNA methylation between treatment timepoints common across samples, we performed paired t-tests, and CpGs were considered significantly differentially methylated before and after treatment if p-adj<0.05 and |FC| > 0.1. In the combination arm 9,460 CpGs were significantly differentially methylated between week 0 and week 12, and 6,940 CpGs were differentially methylated between week 12 and week 25. For the chemotherapy arm, fewer alterations were observed; 649 CpGs between week 0 and week 12, and 311 CpGs between week 12 and week 25 (**Figure 2; Table 2**).

**Figure 2:**
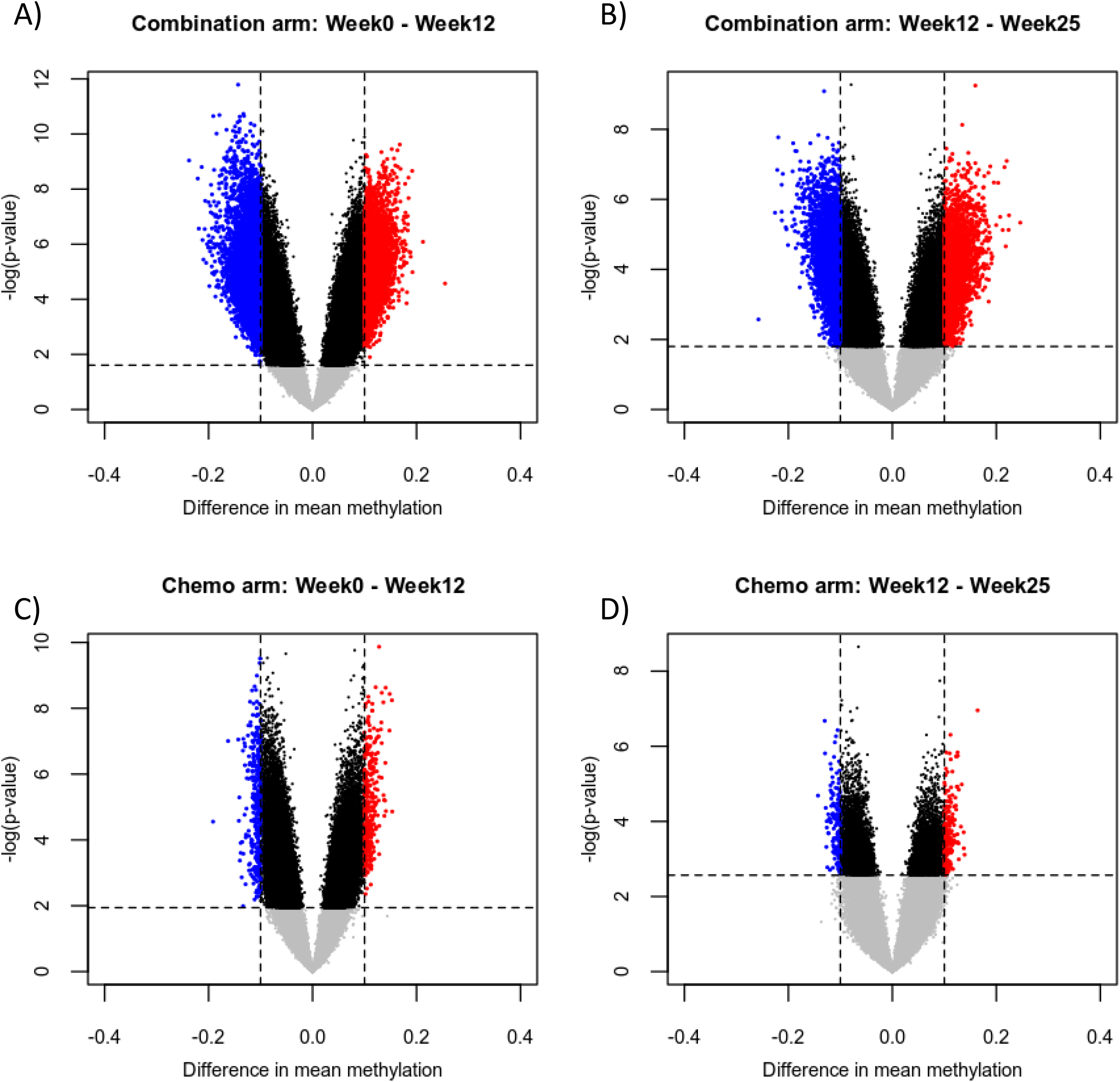
Tumor purity-corrected DNA methylation alterations after treatment with chemotherapy +/- bevacizumab. A) Volcano plots showing differences before and after treatment with chemotherapy (FEC then taxanes) +/- bevacizumab. Matched mean difference in methylation is shown on the x-axis and significance (paired t-test) is shown on the y-axis. CpGs were considered significant if the Benjamini-Hochberg corrected p-values were smaller than 0.05 and the absolute mean difference in methylation was larger than 0.1. Blue dots are CpGs that lose methylation, and red dots are CpGs which gain methylation.

**Table 2:** Changes in DNA methylation during treatment with chemotherapy +/- bevacizumab. The number of significant CpGs with altered DNA methylation levels are shown. Significance was calculated using paired t-tests, and results were considered significant if the Benjamini-Hochberg corrected p-value was smaller than 0.05 and the absolute change in methylation was larger than 0.1. Number of samples in each analysis is shown in parentheses.

|  | Week 0 to week 12 (FEC) | Week 12 to week 25 (Taxanes) |
| --- | --- | --- |
| Combination arm | 9460 (n=36) | 6940 (n=25) |
| Chemo arm | 649 (n=43) | 311 (n=33) |

To assess the function of the genes affected by treatment-induced epigenetic alterations, we mapped CpGs to genes using standard Illumina annotation and performed GSEA of the genes with differential methylation. In general, the affected genes were associated with cell adhesion, differentiation and cell-cell signaling; independent of treatment arm, chemotherapy regimen (FEC or taxanes) and loss/gain of methylation (**Supplementary Figure 2**).

### Integration of DNA methylation and mRNA expression reveals biclusters with distinct biology

We have previously shown that genome-wide associations (*in cis* and *in trans*) between DNA methylation and gene expression (emQTLs) are a powerful approach to identify transcriptional programs based on epigenetic modifications and TF activity, differentially regulated between patients [16, 37]. To identify such epigenetically regulated transcriptional programs involved in response to therapy as well as patient group-specific alterations in DNA methylation, we applied the emQTL approach to delta methylation and delta expression values before and after treatment with FEC +/- Bevacizumab (delta emQTL). The delta emQTL revealed 122,986 significant associations between delta methylation and delta expression (Bonferroni corrected p-value < 0.05), involving 7,921 CpGs and 1,471 genes. Spectral co-clustering was used to identify biclusters of CpGs and genes, representing CpGs and genes with enrichment of significant within-bicluster associations (**Figure 3A**). Two bicluster were chosen based on maximum silhouette scores (**Supplementary Figure 3**) and visual inspection of correlation and anticorrelation between CpGs and genes (**Figure 3A**).

**Figure 3:**
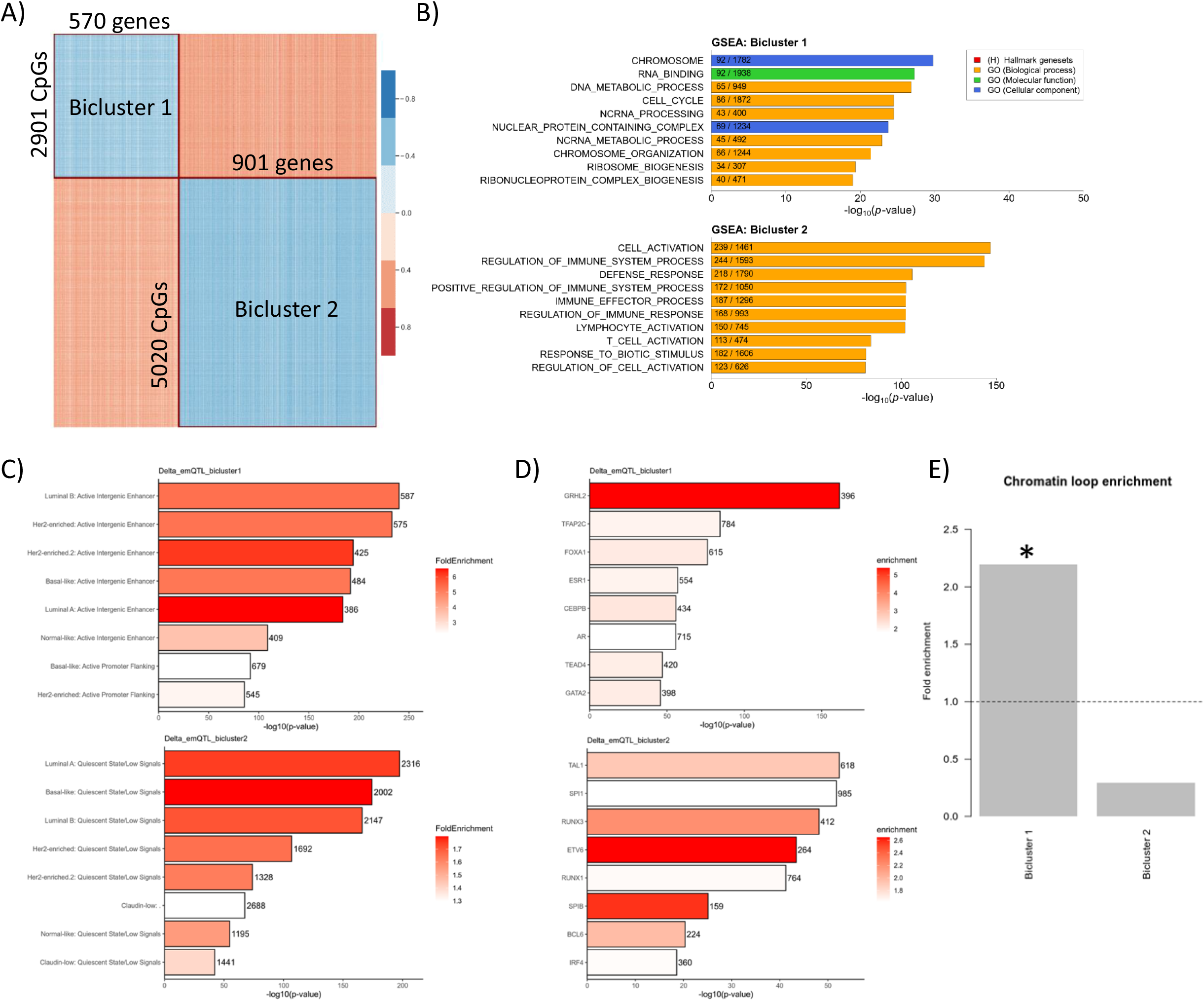
Identification of biclusters of CpGs and genes altered during treatment with FEC +/- bevacizumab. A) Spectral Co-Clustering of correlation coefficients of delta emQTL using k-means identifies two biclusters; heatmap shows ordered rows and columns, where blue represents negative correlation and red represents positive correlation. B) GSEA of genes in the two biclusters. C-D) functional epigenomic enrichment (ChromHMM) analysis of CpGs (C), and transcription factor binding region (UniBind) enrichment analysis (D) of CpGs in Bicluster 1 and Bicluster 2 (upper and lower panels, respectively). P-values (x-axis) are calculated using hypergeometric tests and fill color (C and D) denotes fold enrichment above what is expected by chance. E) Fold enrichment of overlap between CpG-gene pairs in biclusers and chromatin loops determined by ChIA-PET in MCF7.

To assess the putative biological function of each bicluster, we performed GSEA on the gene list from each bicluster. Genes in bicluster 1 were enriched in gene sets associated with chromosomal organization and cell cycle, while Bicluster 2 genes were enriched in gene sets related to immune-related processes (**Figure 3B**). To assess the epigenetic properties of the DNA surrounding the identified CpGs, we utilized ChromHMM segmentation data based on ChIP-seq data of various histone modifications from different breast cancer cell lines covering the PAM50 subtypes. CpGs in Bicluster 1 were significantly enriched in active intergenic enhancer regions of cell lines representing all PAM50 subtypes (**Figure 3C upper panel**), suggesting that methylation of these CpGs may contribute to epigenetic regulation through distal regulatory elements; CpGs in Bicluster 2, conversely, were enriched in regions with quiescent chromatin state in cancer cells (**Figure 3C lower panel**).

We next investigated the TF binding regions surrounding the identified CpGs and TF-DNA interaction data was obtained from UniBind [35]. Bicluster 1 CpGs were found enriched in binding regions of TFs including GRHL2, TFAP2C, FOXA1, ER, CEBPB and NR5A2, TFs known to be important in many cancer-related functions such as proliferation, stemness and EMT (**Figure 3D upper panel**). Bicluster 2 CpGs were enriched in binding regions of TFs such as TAL1, SPI1, RUNX and ETV6, TFs known to be important in the development of the immune system (**Figure 3D lower panel**).

The presence of chromatin loops between enhancers and promoters may suggest epigenetic regulation of genes, and we therefore investigated if the CpGs and genes within the two biclusters were located in chromatin loops of the ER positive breast cancer cell line MCF7 more often than expected by chance. CpGs and genes in Bicluster 1 was significantly enriched in chromatin loops (fold enrichment of 2.2; p-value << 0.05), while CpGs and genes in Bicluster 2 were not enriched (**Figure 3E**). These results further support an epigenetic regulation of genes in Bicluster 1.

### Alterations in DNA methylation with concomitant changes in gene expression associated with treatment response

To assess the molecular alterations during treatment and the relation to treatment response, we performed hierarchical clustering of the delta methylation and delta expression levels of CpGs and genes in the two identified biclusters. For Bicluster 1, patients were divided in three main clusters characterized by strong gain in methylation, moderate gain in methylation and loss of methylation (**Figure 5A**). Tumors in the clusters that showed strong gain of methylation were predominantly ER negative, while tumors in the two other clusters were mainly ER positive. Patients with high gain of methylation showed high rates of pCR (6/15), patients with moderate gain of methylation showed moderate rates of pCR (6/46) while patients with loss of methylation included no patients achieving pCR (0/17; **Figure 5A**). Delta methylation values for Bicluster 2 CpGs showed an inverse pattern compared to Bicluster 1, as the ER negative tumors were enriched in the cluster with strong loss of methylation, while the ER positive patients were divided into two clusters showing either moderate loss of methylation or gain of methylation. The cluster with moderate loss of methylation contained more patients with pCR compared to the cluster with gain of methylation (**Figure 5B**).

To further confirm the association between delta methylation and response to treatment, we averaged the delta methylation levels for all identified CpGs in each bicluster and compared this score against response to treatment (pCR or RCB). Patients achieving pCR or RCB 0 (pCR) or RCB 1 showed increase in delta methylation of Bicluster 1 CpGs and loss of methylation in Bicluster 2 CpGs (p-values < 0.05; **Figure 5C-D and 5F-G**). Further, we performed correlation analyses between the average delta methylation score and the fraction of tumor left after treatment, and we observed a significant correlation for both Bicluster 1- and Bicluster 2-CpGs for all samples and for each arm independently (all p-values < 0.05; **Figure 5E and 5H**). For bicluster 1 CpGs gain of methylation was associated to better response, and for Bicluster 2 CpGs loss of methylation was associated to better response.

The alterations in DNA methylation were accompanied by alterations in gene expression (**Figure 6A-B**). For genes in Bicluster 1 decreased gene expression was associated with better response (**Figure 6C-E**), and for genes in Bicluster 2 increased expression was associated with better response (**Figure 6F-H**).

We performed validation in an independent patient cohort (Doxo/Fumi) where patients with late-stage breast tumors received neoadjuvant treatment with either Doxorubicin or Fluorouracil (5FU) and mitomycin C. In concordance with the observed alterations in DNA methylation from the present patient cohort (NeoAva; **Figure 4**), we observed patient clusters with both gain and loss of DNA methylation for both Bicluster 1 and Bicluster 2 CpGs (**Supplementary Figure 4**). The ER status of tumors was more evenly distributed across the clusters with gain or loss of methylation, perhaps highlighting that the patients in the Doxo/Fumi trial responded less to the treatment [25, 26].

**Figure 4:**
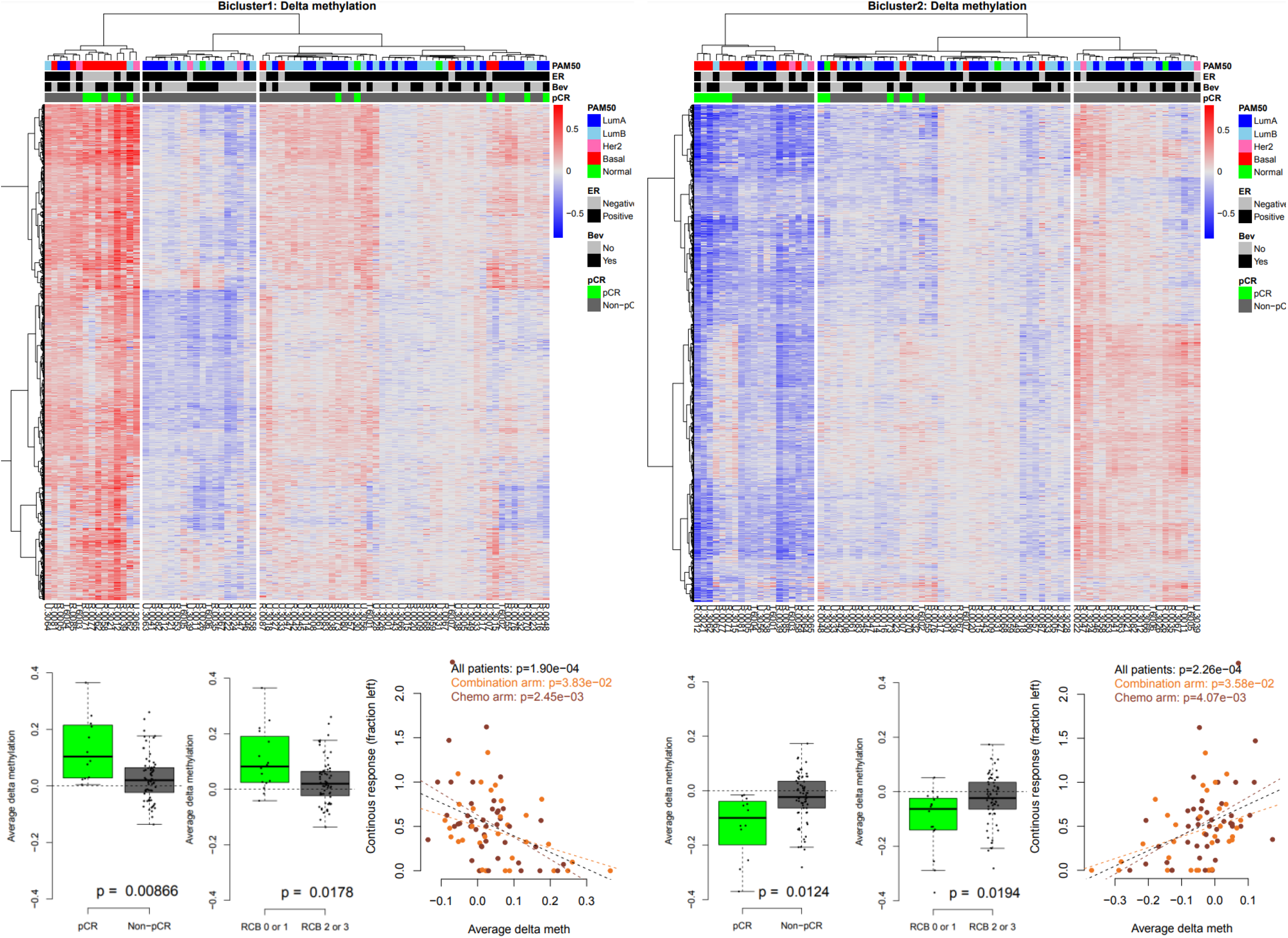
Delta DNA methylation of the identified biclusters after treatment with FEC +/- bevacizumab and association to treatment response. Hierarchical clustering and heatmap of delta methylation values in Bicluster 1 (A) and Bicluster 2 (B); red is gain of methylation and blue is loss of methylation. Three patient clusters were in identified in both dendrograms. Patients (columns) are annotated with PAM50 subtype, ER status, administration of bevacizumab and whether the patient achieved pCR. C-D and F-G) Boxplot showing average delta methylation of Bicluster 1 and 2 CpGs, respectively, plotted against achievement of pCR or RCB. Black dashed line denotes no change in methylation. Statistical significance is calculated using t-test. E and H) Correlation between average delta methylation of Bicluster 1 and 2 CpGs, respectively, and the fraction of tumor left for all patients (black), combination arm (orange) and chemo arm (brown). Statistical significance is calculated using Pearson correlation test.

### Tumor-specific profiles of DNA methylation

Since many of the patients achieved good response to the treatment, it was likely that the observed treatment-induced changes in DNA methylation could be confounded by a higher proportion of non-tumor cells present in the samples taken at week 12 and week 25. To determine alterations in DNA methylation specific to the tumor cells, we developed a method to correct the measured DNA methylation values to represent only tumor cells (see **Materials and methods**).

To identify tumor-specific alterations in DNA methylation between treatment timepoints common across samples, we repeated the paired analyses between timepoints using the tumor purity-corrected DNA methylation levels. In the combination arm, 378 CpGs were differentially methylated between week 0 and week 12 and 5,107 CpGs were differentially methylated between week 12 and week 25. Fewer CpGs were differentially methylated in the chemotherapy arm, 46 and 58 between week 0 and week 12, and week 12 and week 25, respectively (**Table 3** and **Supplementary Figure 5A**). To assess the function of the implicated genes in the tumor cells, we mapped CpGs to genes using standard Illumina annotation and performed GSEA of the genes with differential methylation. From week 0 to week 12 in the combination arm the hypomethylated CpGs were enriched in several gene sets related to adhesion, while the hypermethylated CpGs were enriched in gene sets related to locomotion, migration and cytoskeleton organization (**Supplementary Figure 5B**). From week 12 to week 25 in the combination arm both hypomethylated and hypermethylated CpGs were enriched in gene sets related to adhesion and locomotion (**Supplementary Figure 5B**). In the chemotherapy arm, too few CpGs were differentially methylated to perform GSEA.

**Table 3:**
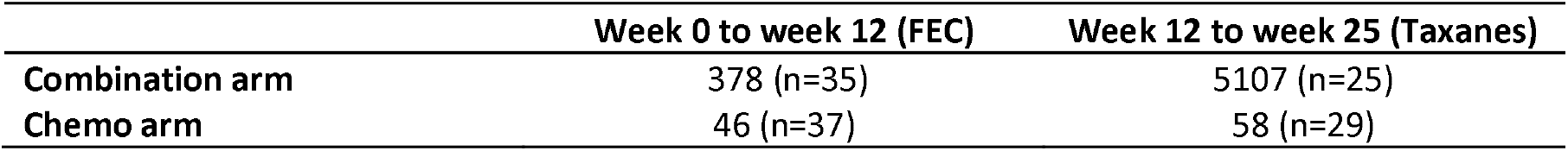
Changes in tumor purity-corrected DNA methylation during treatment with chemotherapy +/- bevacizumab. The number of significant CpGs with altered DNA methylation levels are shown. Significance was calculated using paired t-tests, and results were considered significant if the Benjamini-Hochberg corrected p-value was smaller than 0.05 and the absolute change in methylation was larger than 0.1. Number of samples in each analysis is shown in parentheses.

|  | Week 0 to week 12 (FEC) | Week 12 to week 25 (Taxanes) |
| --- | --- | --- |
| Combination arm | 378 (n=35) | 5107 (n=25) |
| Chemo arm | 46 (n=37) | 58 (n=29) |

To assess whether the delta emQTL results (**Figure 5**) were influenced by lower proportion of tumor cells in the treated biopsies, we performed hierarchal clustering also on the tumor purity-adjusted delta DNA methylation levels. Delta methylation levels of Bicluster 1 CpGs showed concordant patterns (compared to non-adjusted data) across tumors, with two clusters showing gain of methylation and one showing loss of methylation (**Supplementary Figure 6A**) suggesting that the altered methylation of these CpGs were likely caused by alterations in the tumor cells or altered composition of tumor subclones. Delta methylation levels of Bicluster 2 CpGs also showed loss of methylation for the ER negative tumors, but the pattern for ER positive tumors was less clear (**Supplementary Figure 6B**), again implying that methylation levels of these CpGs were more influenced by non-tumor cells such as immune cells. When comparing average tumor purity-adjusted delta methylation to response, we observed that the differences in average methylation was still (borderline) significant for Bicluster 1 CpGs (pCR p-value=0.0518; RCB p-value=0.0323; **Supplementary Figure 6C-D)**, but not significant for CpGs in Bicluster 2 (pCR p-value=0.168; RCB p-value=0.0812; **Supplementary Figure 6E-F**)

**Figure 5:**
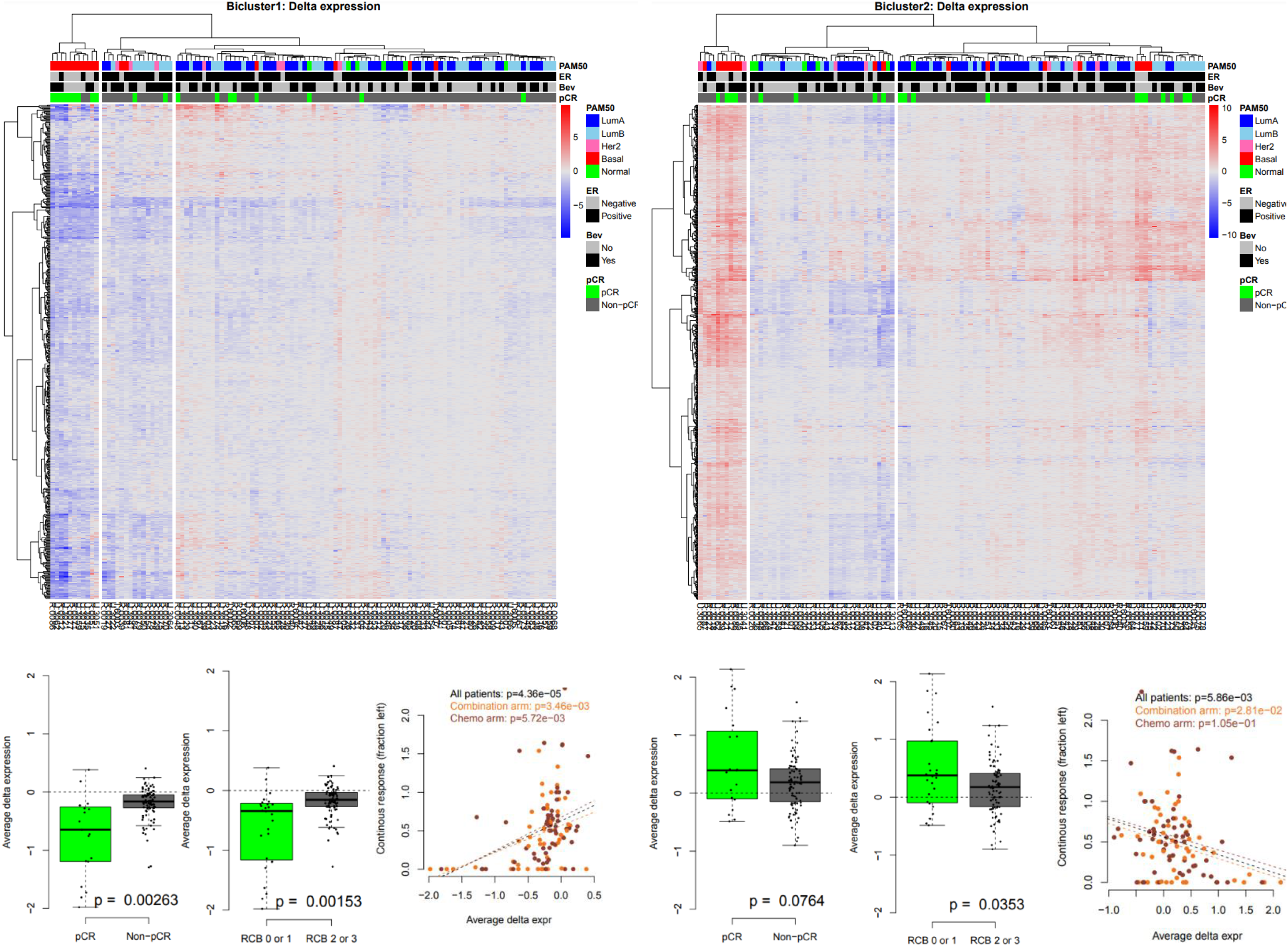
Delta gene expression of the identified biclusters after treatment with FEC +/- bevacizumab and association to treatment response. A-B) Hierarchical clustering and heatmap of delta expression values; red is gain of expression and blue is loss of expression. Three patient clusters were in identified in both dendrograms. Patients (columns) are annotated with PAM50 subtype, ER status, administration of bevacizumab and whether the patient achieved pCR. C-D and F-G) Boxplot showing average delta expression of Bicluster 1 and 2 genes, respectively, plotted against achievement of pCR or RCB. Black dashed line denotes no change in expression. Statistical significance is calculated using t-test. E and H) Correlation between average delta expression of Bicluster 1 and 2 genes, respectively, and fraction of tumor left for all patients (black), combination arm (orange) and chemo arm (brown). Statistical significance is calculated using Pearson correlation test.

To confirm that the predictive signature was not confounded by tumor purity, we applied the stable model (**Table 1**) on tumor purity-adjusted DNA methylation data and we achieved a perfect prediction performance (AUC=1), in concordance with the results obtained using non-adjusted DNA methylation data.

## Discussion

Here we present DNA methylation profiles of fresh frozen tumor biopsies taken before, during and after treatment from patients receiving neoadjuvant chemotherapy with or without bevacizumab. We report that DNA methylation at enhancer CpGs related to cell cycle, estrogen response and immune infiltration can predict response to the combination of chemotherapy and bevacizumab, and we validate this in an independent patient cohort from a similar clinical trial [24]. The prediction of treatment response is further improved when combining the DNA methylation score with the previously published ViRP proteomic score [12]. We have previously reported that administration of bevacizumab in addition to chemotherapy may confer improved response for patients [11], and here we show that tumors receiving the combination treatment underwent more extensive epigenetic alterations than tumors receiving only chemotherapy both in bulk and when corrected for tumor purity. We then performed integrative emQTL analysis on alterations in DNA methylation and gene expression levels, showing that the epigenetic alterations that occur during treatment are different between responders and non-responders and that these differences may be explained by the proliferation-EMT axis through the activity of GRHL2 and other TFs.

The use of Bevacizumab plus chemotherapy as first-line treatment in patients with locally advanced or metastatic breast cancer has shown improved response and clinical benefit in the present study (NeoAva) [11] and others [6, 9, 10, 38, 39]. Here, we present a predictive biomarker for selection of patients with ER positive HER2 negative tumors that have a high probability of improved response by addition of bevacizumab. Since ER positive and ER negative tumors are different with regards to drivers of proliferation, it is an advantage that the predictive signature is trained and validated on ER positive tumors specifically. Taken together, these results illustrate the clinical benefit of bevacizumab if administered to the correct patients.

Preselection of CpGs for identification of the predictive signature was performed based on the results in Ankill et al. [17], where we show that 44,263 CpGs capture important biological variation in breast cancer involving estrogen signaling, non-estrogen-controlled regulation of proliferation, immune infiltration and fibroblast infiltration. The majority of CpGs in the identified predictive signature were associated to regulation of the cell cycle, showing that epigenetic marks related to proliferation can be predictive of response, which is in concordance with previous studies [15, 21].

The delta emQTL approach has the advantage of identifying concomitant alteration in DNA methylation and gene expression that are different between patients, which allowed us to identify CpGs and genes whose alterations were associated to response. We identified two biclusters of CpGs and genes with negative within-bicluster correlations, one with enrichment in enhancers and TFs important in functions such as proliferation, stemness and EMT, and one with enrichment in immune-related TFs, both of which were (oppositely) associated with response. We validated the profiles of alterations in DNA methylation during chemotherapy in an independent patient cohort treated with doxorubicin or a combination of 5-FU and mitomycin, but we did not validate the association to response; the lack of validation to response may be due to that the two patient cohorts were collected at different time periods, and that the tumors in the validation cohort were larger, and the response rates were a lot poorer overall.

The association between treatment response and alteration of methylation in enhancer CpGs enriched in GRHL2 binding regions with concomitant changes of proliferation-related genes (bicluster 1) suggests that proliferative cells with high activity of GRHL2 are killed by the treatment leading to an observed gain of methylation and loss of expression (**Figure 5A and 6A**). GRHL2 (Grainyhead-like-2) is known to promote proliferation and suppress EMT by repressing ZEB1 expression and inhibiting TGF-β signaling [40], and to suppress the emergence of a stem cell phenotype [41]. We observe that gain of GRHL2 activity through loss of enhancer methylation with following increase in proliferation-related genes is characteristic of non-responders, suggesting that non-responding tumor cells can gain proliferative potential during treatment stress. For bicluster 2, the observed alterations in methylation and expression suggest that eradication of tumor cells lead to a relative increase in non-tumor cells (e.g., immune cells) in the tumors that respond well to treatment, which in turn is reflected in loss of methylation at enhancers carrying TF binding regions of TFs related to immune development and increase in expression of immune-related genes.

In this study we present a method for adjusting DNA methylation levels to reflect tumor cells only (tumor purity-adjusted DNA methylation), when tumor purity estimates are available. In our results, analyses with tumor purity-adjusted data did not lead to new biological interpretation. We observed fewer differentially methylated CpGs, but the genes affected were enriched in the same biological functions (i.e., adhesion, locomotion etc.; **Supplementary Figures 2 and 5**). The method for tumor purity-corrected DNA methylation uses a linear regression approach where values are replaced with the residual of the regression model. This approach is commonly used when correcting gene expression values for unwanted covariates such as RIN value. In this case, the approach assumes that the methylation level of each CpG is equal in all non-tumor cells, and that deviations from “expected” (i.e., the regression line) is attributed to tumor cells. Since DNA methylation alterations in breast tumors are large and frequent, this assumption is reasonable, but not true for all CpGs. Nonetheless, the method is useful when studying tumors where tumor purity is an important and unwanted confounding factor.

## Conclusion

Here we report that DNA methylation at enhancer CpGs related to cell cycle regulation can predict response to the treatment with chemotherapy and bevacizumab for ER positive patients and we validate this observation in an independent patient cohort. When combining the DNA methylation score with a previously reported proteomic score, the prediction further improved. We also show that tumors receiving the combination treatment underwent more extensive epigenetic alterations than tumors receiving only chemotherapy. Finally, we perform the integrative emQTL analysis on alterations in DNA methylation and gene expression levels, showing that the epigenetic alterations occurring during treatment differs between responders and non-responders and that these differences may be explained by the proliferation-EMT axis through the activity of the transcription factor GRHL2. Taken together, these results illustrate the importance of predictive markers to identify patients with clinical benefit of the addition of bevacizumab to chemotherapy.

## Acknowledgments

The contribution to the study from all the participating patients is greatly acknowledged.

