## Supplementary figures for "An integrated ‘omics approach highlights the role of epigenetic events to explain and predict response to neoadjuvant chemotherapy and bevacizumab"

### Slide 1
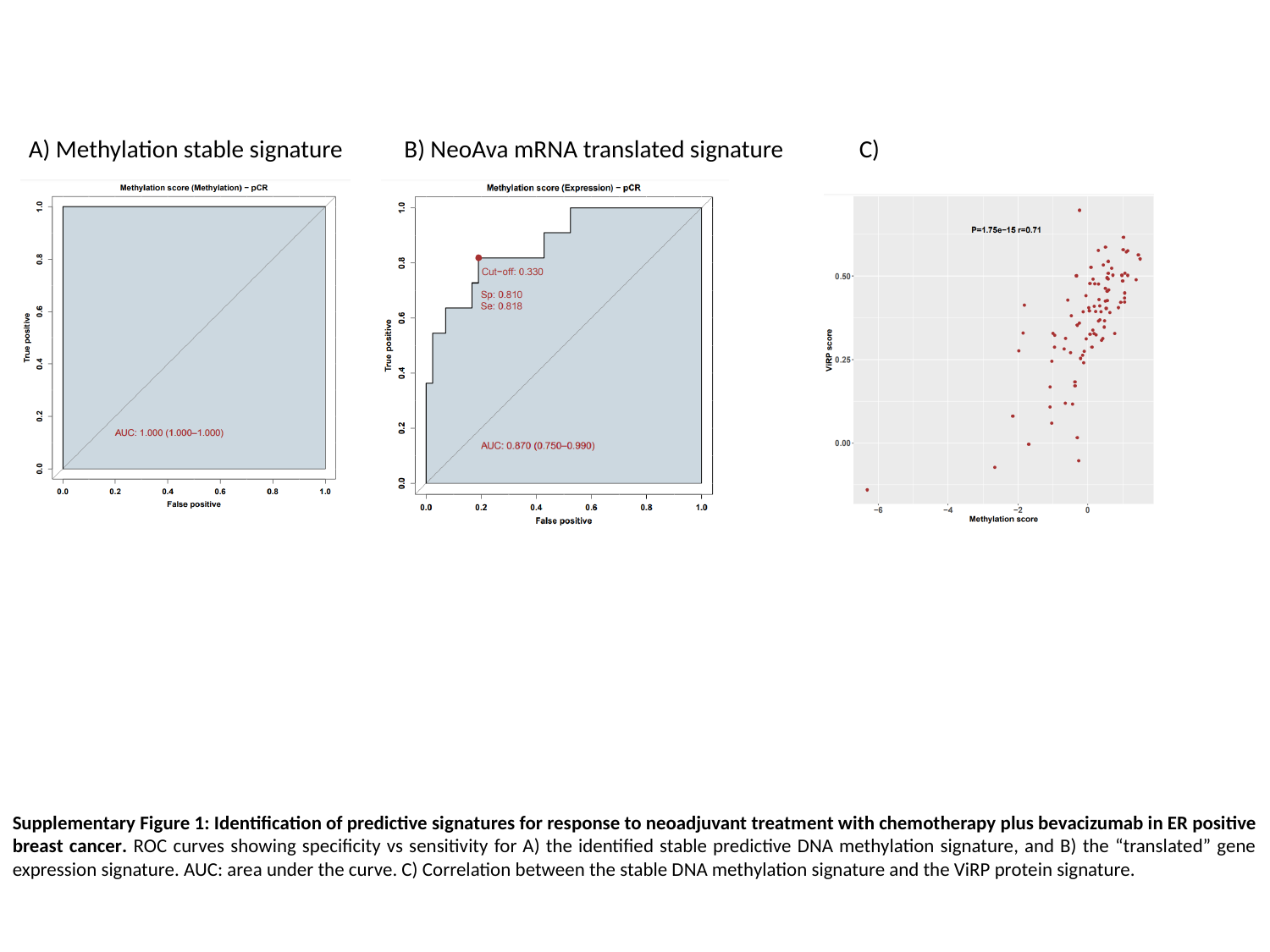

A) Methylation stable signature
B) NeoAva mRNA translated signature
C)
Supplementary Figure 1: Identification of predictive signatures for response to neoadjuvant treatment with chemotherapy plus bevacizumab in ER positive breast cancer. ROC curves showing specificity vs sensitivity for A) the identified stable predictive DNA methylation signature, and B) the “translated” gene expression signature. AUC: area under the curve. C) Correlation between the stable DNA methylation signature and the ViRP protein signature.

### Slide 2
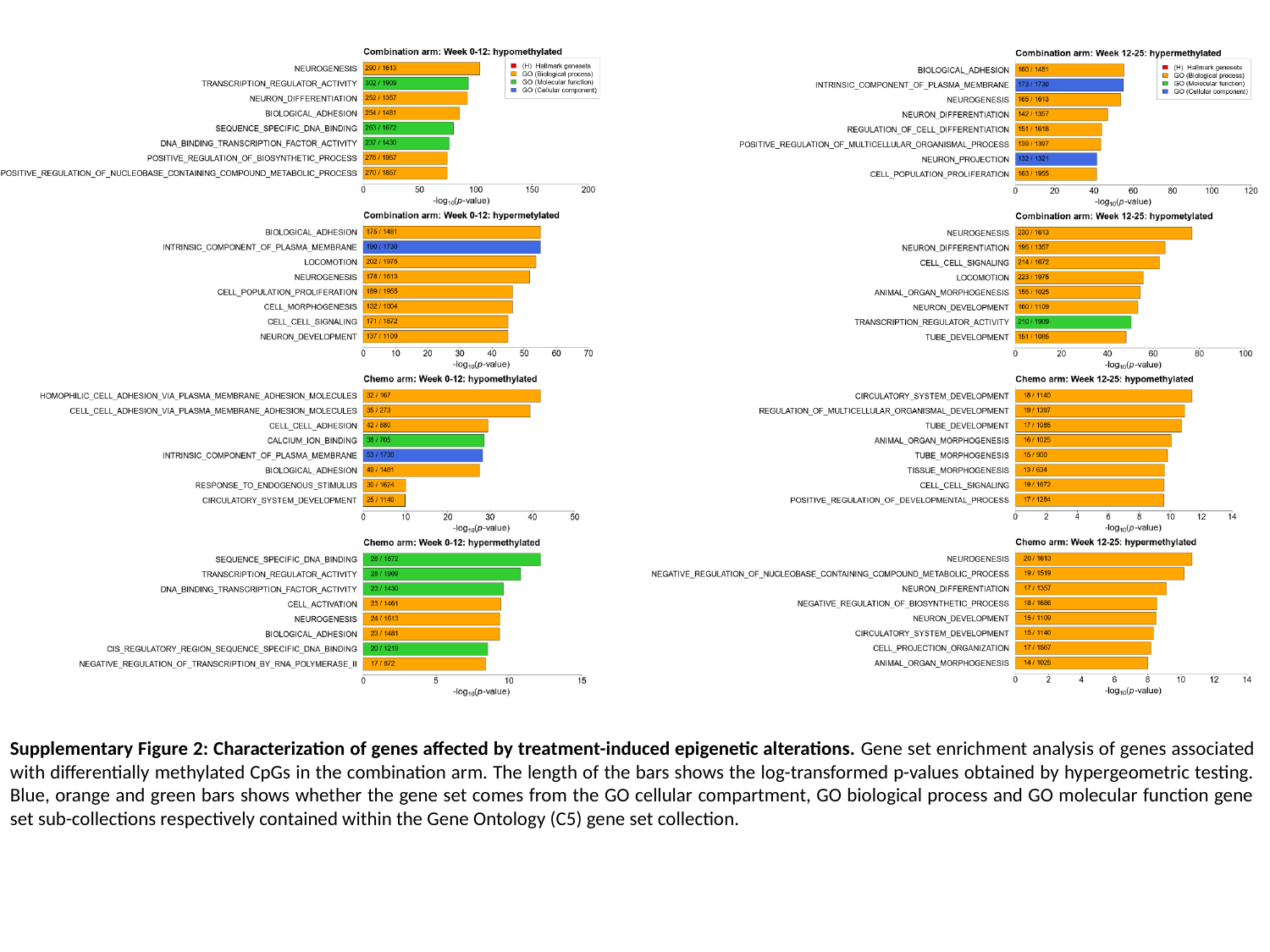

Supplementary Figure 2: Characterization of genes affected by treatment-induced epigenetic alterations. Gene set enrichment analysis of genes associated with differentially methylated CpGs in the combination arm. The length of the bars shows the log-transformed p-values obtained by hypergeometric testing. Blue, orange and green bars shows whether the gene set comes from the GO cellular compartment, GO biological process and GO molecular function gene set sub-collections respectively contained within the Gene Ontology (C5) gene set collection.

### Slide 3
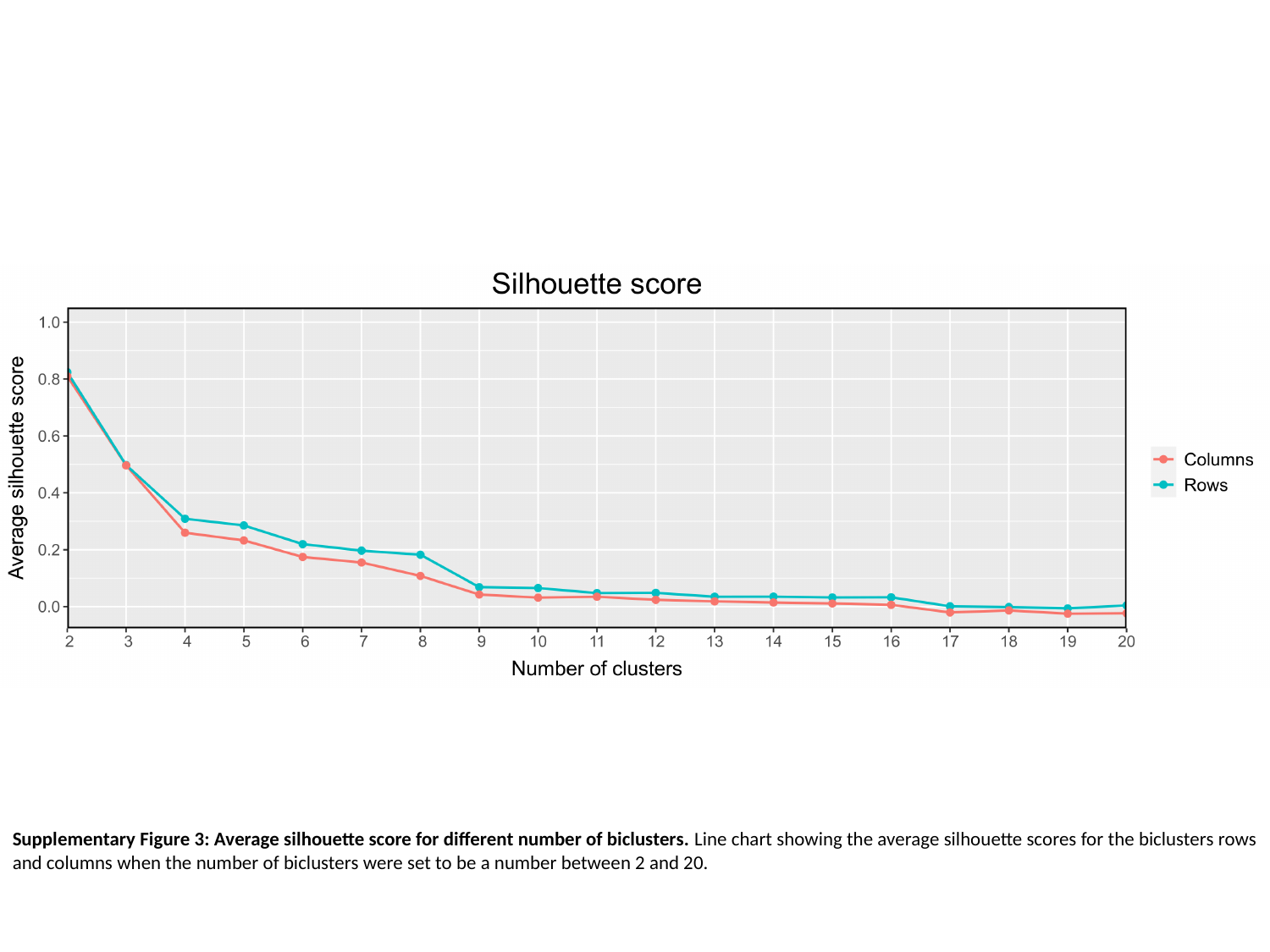

Supplementary Figure 3: Average silhouette score for different number of biclusters. Line chart showing the average silhouette scores for the biclusters rows and columns when the number of biclusters were set to be a number between 2 and 20.

### Slide 4
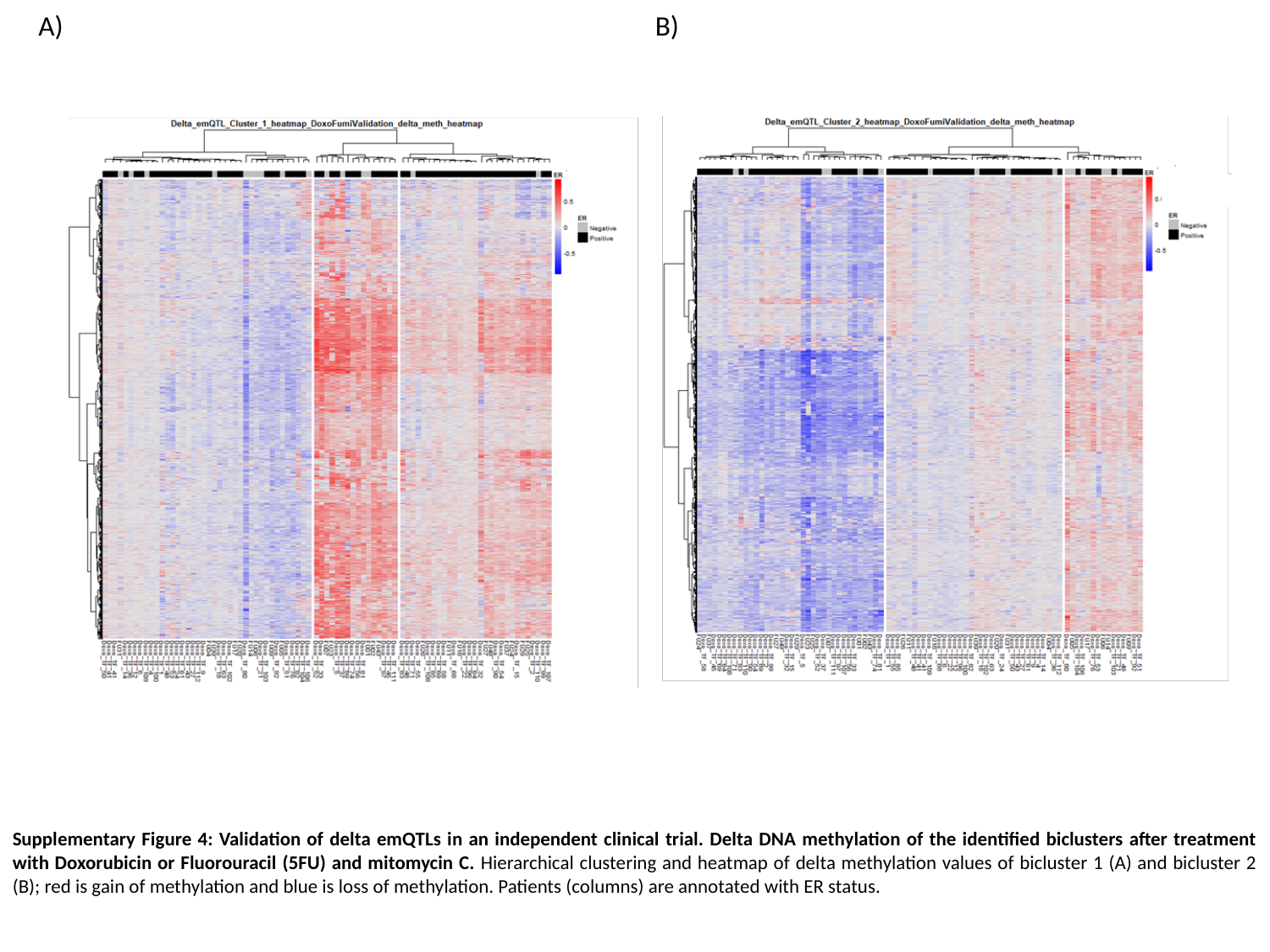

A)
B)
Supplementary Figure 4: Validation of delta emQTLs in an independent clinical trial. Delta DNA methylation of the identified biclusters after treatment with Doxorubicin or Fluorouracil (5FU) and mitomycin C. Hierarchical clustering and heatmap of delta methylation values of bicluster 1 (A) and bicluster 2 (B); red is gain of methylation and blue is loss of methylation. Patients (columns) are annotated with ER status.

### Slide 5
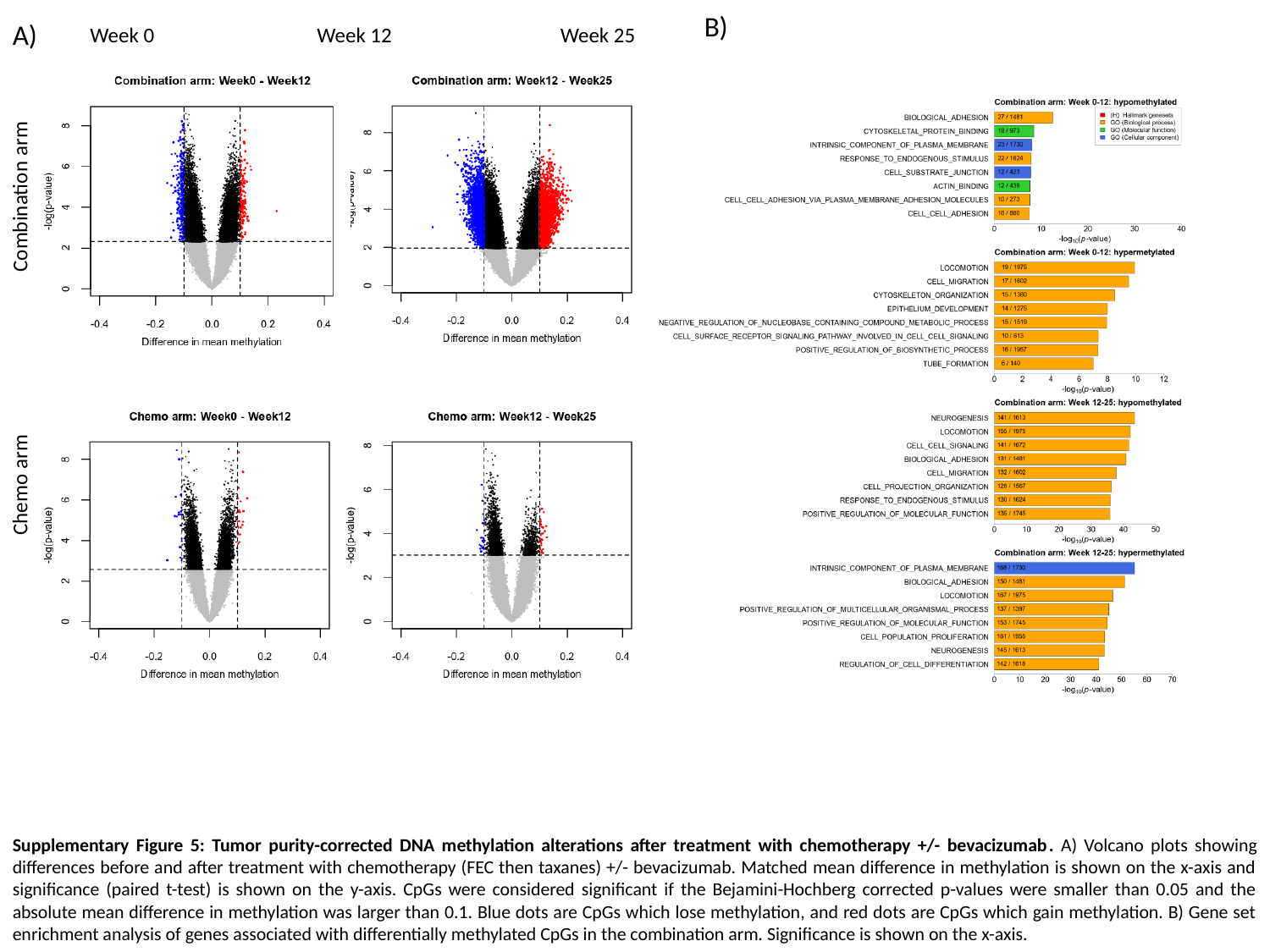

B)
A)
Week 0
Week 12
Week 25
Combination arm
Chemo arm
Supplementary Figure 5: Tumor purity-corrected DNA methylation alterations after treatment with chemotherapy +/- bevacizumab. A) Volcano plots showing differences before and after treatment with chemotherapy (FEC then taxanes) +/- bevacizumab. Matched mean difference in methylation is shown on the x-axis and significance (paired t-test) is shown on the y-axis. CpGs were considered significant if the Bejamini-Hochberg corrected p-values were smaller than 0.05 and the absolute mean difference in methylation was larger than 0.1. Blue dots are CpGs which lose methylation, and red dots are CpGs which gain methylation. B) Gene set enrichment analysis of genes associated with differentially methylated CpGs in the combination arm. Significance is shown on the x-axis.

### Slide 6
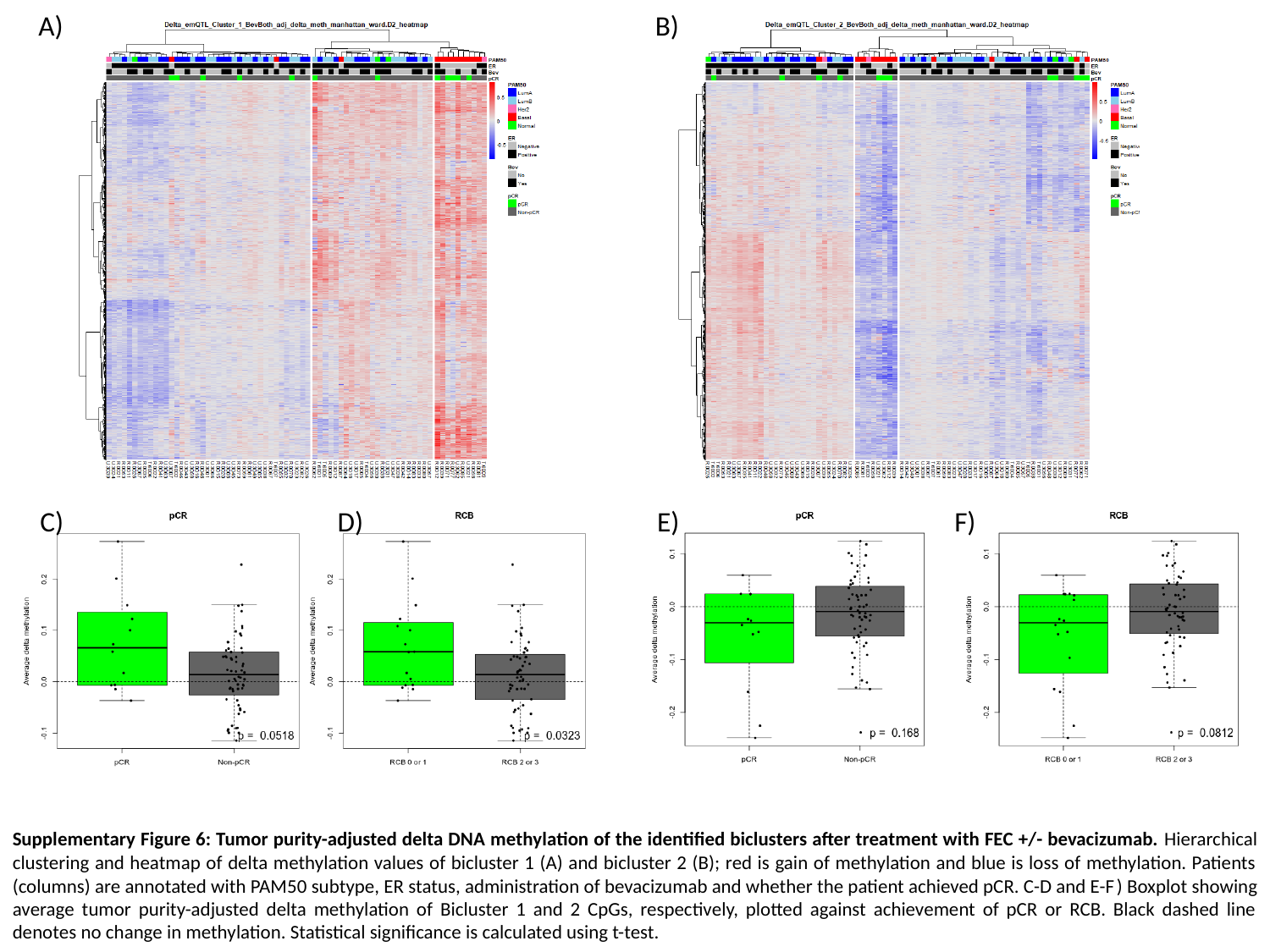

A)
B)
C)
D)
E)
F)
Supplementary Figure 6: Tumor purity-adjusted delta DNA methylation of the identified biclusters after treatment with FEC +/- bevacizumab. Hierarchical clustering and heatmap of delta methylation values of bicluster 1 (A) and bicluster 2 (B); red is gain of methylation and blue is loss of methylation. Patients (columns) are annotated with PAM50 subtype, ER status, administration of bevacizumab and whether the patient achieved pCR. C-D and E-F) Boxplot showing average tumor purity-adjusted delta methylation of Bicluster 1 and 2 CpGs, respectively, plotted against achievement of pCR or RCB. Black dashed line denotes no change in methylation. Statistical significance is calculated using t-test.
